# Super-human cell death detection with biomarker-optimized neural networks

**DOI:** 10.1101/2020.08.04.237032

**Authors:** Jeremy W. Linsley, Drew A. Linsley, Josh Lamstein, Gennadi Ryan, Kevan Shah, Nicholas A. Castello, Viral Oza, Jaslin Kalra, Shijie Wang, Zachary Tokuno, Ashkan Javaherian, Thomas Serre, Steven Finkbeiner

## Abstract

Cell death is an essential process in biology that must be accounted for in live microscopy experiments. Nevertheless, cell death is difficult to detect without perturbing experiments with stains, dyes or biosensors that can bias experimental outcomes, lead to inconsistent results, and reduce the number of processes that can be simultaneously labelled. These additional steps also make live microscopy difficult to scale for high-throughput screening because of the cost, labor, and analysis they entail. We address this fundamental limitation of live microscopy with biomarker-optimized convolutional neural networks (BO-CNN): computer vision models trained with a ground truth biosensor that detect live cells with superhuman, 96% accuracy more than 100 times faster than previous methods. Our models learn to identify important morphological characteristics associated with cell vitality without human input or additional perturbations, and to generalize to other imaging modalities and cell types for which they have no specialized training. We demonstrate that we can interpret decisions from BO-CNN models to gain biological insight into the patterns they use to achieve superhuman accuracy. The BO-CNN approach is broadly useful for live microscopy, and affords a powerful new paradigm for advancing the state of high-throughput imaging in a variety of contexts.

## Introduction

Observing dynamic biological processes over time with live fluorescence microscopy has played an essential role in establishing our understanding of fundamental cell biology. Nevertheless, image analysis is often a rate-limiting step for scientific discovery^1, 2^. It frequently requires labor-intensive manual curation, annotation, and quantification of specific visual features, which dramatically reduces throughput and potentially introduces human bias. The advent of high-throughput automated microscopy has increased the rate of data generation without solving critical problems affecting the rate and quality of data analysis^3^. Without better approaches that overcome the inefficiencies introduced by manual image analysis, the impact of imaging on scientific discovery will be limited.

A popular alternative to manual curation is semi-automated image analysis, available through open-source platforms such as ImageJ/National Institutes of Health (NIH)^4^ and CellProfiler^5^. This approach relies on feedback loops between manual curation and basic statistical models to incrementally build better computer vision models for rapidly phenotyping samples^6^. However, the resulting models rely on visual features that are highly tuned to the specific experimental conditions they were trained on, meaning that new models must be built for every new experiment. These approaches are also ultimately limited by the precision and accuracy with which human curators can score and annotate images. Variability in curator precision, combined with ambiguities present in biological imaging due to technical artifacts, often lead to a noisy ground truth, limiting the success of attempts towards automation. The recent explosion of computer vision models leveraging deep learning has led to more accurate, versatile, and efficient algorithms for cellular image analysis^7^.

Convolutional neural networks (CNNs) in particular have been responsible for massive improvements over classic computer vision in segmentation within electron microscopy^8-10^ or microscopy^11^, automated phenotypic biomarker analysis in live cell microfluidics^12^, subcellular protein localization^13^ and fluorescent image segmentation^14^, and identification of cell lineages^15^. For example, CNNs have been found to be able to classify hematoxylin- and eosin-stained histopathology slides with super-human accuracy^16^, and these models can sometimes generalize between datasets with minimal or no additional labeled data^17^. Remarkably, CNNs have shown promise in detecting visual features within images that are undetectable by humans, suggesting that they can be harnessed to drive scientific discovery^18^. Nevertheless, a major factor in achieving such strong performance for CNNs is training with large labeled data sets, which can require massive amounts of human curation and/or annotation^19, 20^. Thus, it is critical to develop methods that can generate datasets which are sufficiently large to train CNNs, but with minimal human curation.

Cell death is a ubiquitous phenomenon within live imaging data requiring substantial attention in image analysis. The physiological properties of live and dead cells are fundamentally different, and it is critical to distinguish live from dead cells within a population. In some cases, cell death is noise that must be filtered from signal coming from live cells, while in others death is a biological phenotype that must be analyzed, for instance in studies of development^21^, cancer^22^, ageing^23^ and disease^24^.

In cellular models of neurodegenerative disease, neuronal death is frequently used as a disease-relevant phenotype^25–27^, and its analysis as a tool for the discovery of covariates of neurodegeneration^28–30^. Nevertheless, detection of neuronal death can be challenging, especially in live imaging studies, due to the dearth of acute, time-resolved biosensors capable of detecting death. A plethora of cell death indicators, dyes, and stains have been developed and described, yet application of these reagents can introduce artificial toxicity into an experiment, they are often difficult to quantify and display significant batch-to-batch variability, and they tend to report very late stages of death after significant decay has already occurred^27^. More recently, we established a novel family of genetically encoded death indicators (GEDI) that acutely mark a stage at which neurons are irreversibly committed to die^31^. Our approach uses a robust signal that standardizes the measurement across experiments and is amenable to high-throughput analysis. However, this approach cannot be applied to cells that do not express the GEDI reporter. Moreover, the GEDI construct emits in two fluorescent channels, which restricts its use in cells co-expressing fluorescent biosensors in overlapping emission spectra.

Here we address the extant issues in detecting neuronal death in microscopy by developing a novel quantitative microscopy pipeline that automatically generates GEDI-curated data to train a CNN without human input. The resulting GEDI-CNN is capable of detecting neuronal death from images of morphology alone, alleviating the need for any additional use of GEDI in experiments. The GEDI-CNN achieves super-human accuracy and speed in identifying neuronal cell death in high-throughput live-imaging data sets.

Through systematic analysis of a trained GEDI-CNN, we find that it learns to detect death in neurons by locating morphology that has classically been linked to death, despite receiving no explicit supervision towards these features. We also show this model generalizes to images captured with different parameters or displays of neurons and cell types from different species without additional training, including to studies of neuronal death in cells derived from patients with neurodegenerative disease. These data demonstrate a versatile and promising new application of CNN models for aiding analysis and discovery in biological studies using live microscopy data, as well as a novel approach for optimizing CNNs using biomarker signals directly.

## Results

### An automated pipeline for optimizing a GEDI-based classifier for neuronal death

Previously we showed that the GEDI biosensor can detect irreversibly elevated intracellular Ca^2+^ levels to quantitatively and definitively classify neurons as live or dead^31^ (Figure 1A-C). In contrast to other live/dead indicators, GEDI is constitutively expressed in cells to quantitatively and irreversibly report death only when a cell has reached a level of calcium not found in live cells. Prior to the GEDI biosensor, human curation of neuronal morphology was often deployed for detection of death in longitudinal imaging data sets, and was considered the “gold standard”^27, 32^. It has been long recognized that the transition from life to death closely tracks with changes in a neuron’s morphology that occur as it degenerates^33^. However human curation based on morphology is subjective, relies on subjective and ill-defined interpretation of features in an image, and thus can be both inaccurate and imprecise^31, 34^. Furthermore, in cases Figure 1 where human curation is incorrect, it remains unknown whether images of cell morphology convey enough information to indicate an irreversible commitment to death, or whether curators incorrectly identify and weigh features within the dying cells.

**Figure 1.**
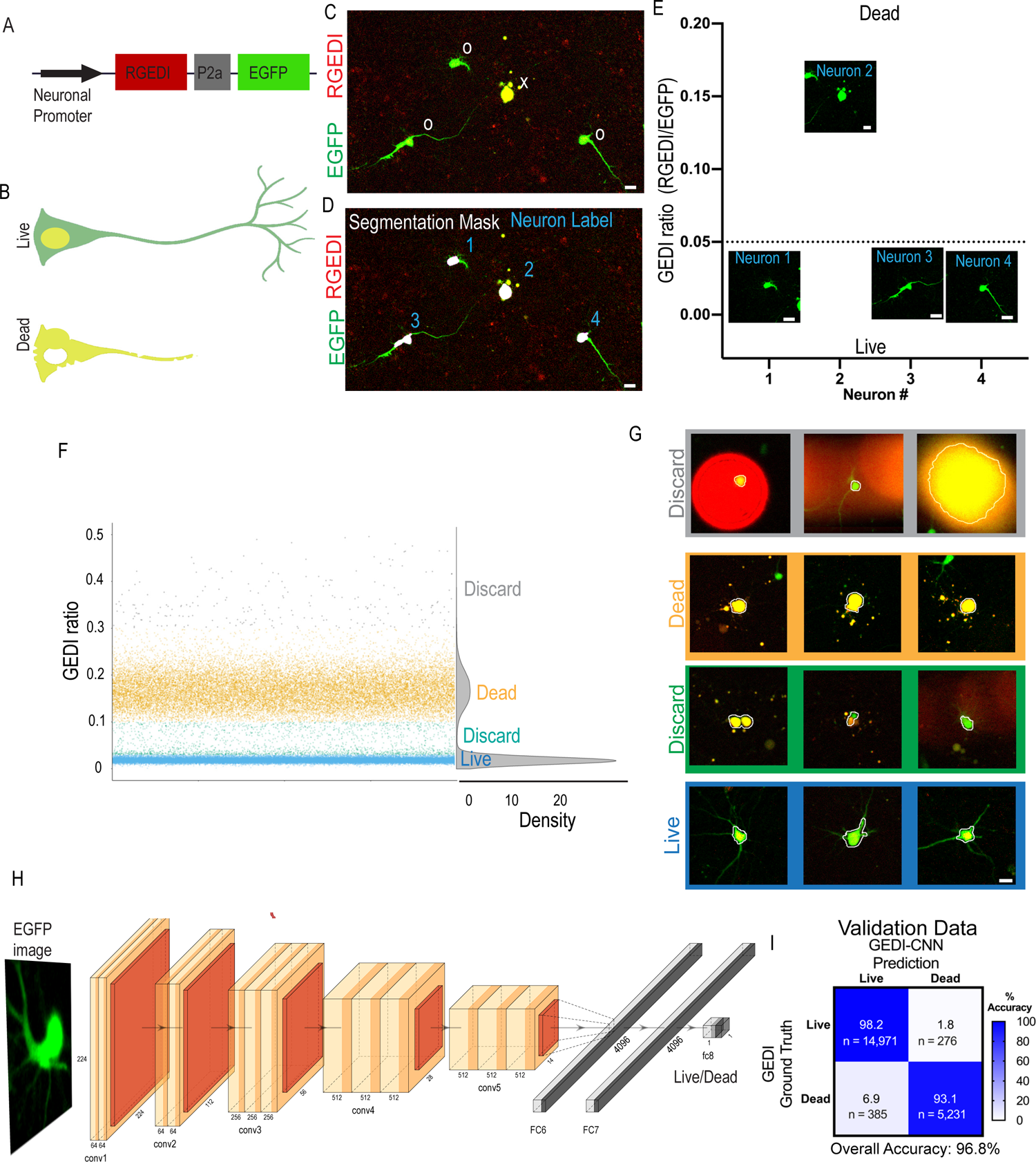
GEDI signal as a ground truth for training a live/dead classifier CNN from morphology. A) The GEDI biosensor expression plasmid contains a neuron-specific promoter driving expression of a red fluorescent RGEDI protein, a P2a “cleavable peptide”, and an EGFP protein. Normalizing the RGEDI signal to the EGFP signal (GEDI Ratio) at a single cell level provides a ratiometric measure of a “death” signal that is largely independent of cell-to-cell variation in transfection efficiency and expression. B) Schematic overlay of green and red channels illustrating the GEDI sensor’s color change in live neurons (top) and dead neurons (bottom). Live neurons typically contain basal RGEDI signal in the nucleus and in the perinuclear region near intracellular organelles with high Ca^2+^ ^31^. C) Representative red and green channel overlay of neurons expressing GEDI, showing one dead (x) and three live (0) neurons. D) Segmentation of image in (C) for objects above a specific size and intensity identifies the soma of each neuron (segmentation masks), which is given a unique identifier label (1–4). E) Ratio of GEDI to EGFP fluorescence (GEDI ratio) in neurons from (D). A ratio above the GEDI threshold (dotted line) indicates an irreversible increase in GEDI signal associated with neuronal death. Cropped EGFP images are plotted at the level of their associated GEDI ratio. (F, G) Generation of GEDI-CNN training data sets from images of individual cells. GEDI ratios from images of each cell (G) were used to create training examples of live and dead cells. Cells with intermediate or extremely high GEDI ratios were discarded to eliminate ambiguity during the training process. Automated cell segmentation boundaries are overlaid in white. H) Architecture of GEDI-CNN model based off of VGG16 architecture. I) Accuracy confusion matrix for live/dead classification. Scale bar = 20μm.

Because the GEDI biosensor gives a readout of a specific physiological marker of a cell’s live/dead state, it provides superior fidelity in differentiating between live and dead cells^31^. We hypothesized that the GEDI biosensor signal could be leveraged to recognize consistent morphological features of live and dying cells and help improve classification in datasets where GEDI is not present. Additionally, we sought to further understand how closely morphology features track with death and whether early morphological features of dying cells could be more precisely identified and defined. To this end, we optimized convolutional neural networks (CNNs) with the GEDI biosensor signal.

Neurons transfected with a GEDI construct co-express the red fluorescent protein (RGEDI) and green fluorescent EGFP (morphology label) in a one-to one ratio due to their fusion by a porcine teschovirus-1 2a (P2a) “self-cleaving” peptide^31^ (Figures 1A-B). RGEDI fluorescence increases when the Ca^2+^ level within a neuron reaches a level indicative of an irreversible commitment to death. We normalize for cell-to-cell variations in the amount of the plasmid transfected by dividing the RGEDI fluorescence by the fluorescence of EGFP, which is constitutively expressed from the same plasmid, to give the ratiometric GEDI ratio (Figure 1C). Starting from our previously described time series of GEDI-labelled primary cortical neurons from rats (31), we segmented the EGFP-labeled soma of each neuron with an intensity threshold and minimum and maximum size filter, as previously described^35^. Using the intensity weighted-centroid from each segmentation mask, images of each neuron were automatically cropped and sorted based on their GEDI ratio as live or dead (Figure 1D–E). Plots of the GEDI ratio from the first time point of the data set showed that the values from most neurons clustered into two groups corresponding to either live or dead cells (Figure 1F). A minority of neurons had GEDI ratios outside the main distribution of live and dead GEDI signals, typically due to segmentation errors (Figure 1G), and these were discarded from the training set. Cropped images from a total of 53,638 live and 22,533 dead neurons from the first timepoint of an imaging series, both with and without background subtracted, were sorted to generate a data set for training computer vision models. We selected a widely-used CNN called VGG16, which was initialized with weights from training on the large-scale ImageNet dataset of natural images^36^. We refer to this model as GEDI-CNN (Figure 1H, Supplemental Figure 1). In validation on held-out data from the first timepoint of the imaging series, the GEDI-CNN classified dead cells from the EGFP channel images with 96.8% accuracy (0.96 Area Under the ROC curve (AUC)) in (Figure 1I). The GEDI-CNN did not have diminished capacity in classifying neurons in the validation set with GEDI ratios that were discarded from the training data because their GEDI signal fell in regions outside the main distribution (Supplemental Figure 1), suggesting the removal of discarded neurons during training did not distort the model.

However, this did not diminish the GEDI-CNN’s capacity to classify neurons in the validation set, suggesting the removal of discarded these neurons during training did not distort the model. These data indicate that direct utilization of a biomarker signal without human curation can generate a highly accurate biomarker-optimized CNN capable of automated classification of microscopy images.

### Applying GEDI-CNN to longitudinal imaging of individual neurons

Live cell imaging over multiple time points exacerbates the problems with classification of dead cells by human curators because each time point multiplies the number of images that must be classified. Therefore, GEDI-CNN curation could be especially advantageous for cell-imaging time series. As the GEDI-CNN model was trained only with the first time point of our imaging series, we next tested its ability to classify individual neurons at later time points. Across 478,987 neurons tested (Supplemental Figure 1), the GEDI-CNN was 96.4% accurate (0.92 AUC) in detecting dead neurons, with little deviation across 20 different imaging plate batches (range: 88.4%-97.4%) and across 46 different days of imaging (range: 87.8%-98.1%) (Figure 2A). The GEDI-CNN was slightly better at correctly classifying live neurons than dead neurons, but was still able to successfully classify datasets with imbalanced numbers of live vs. dead cells (Figure 2B, C). The focal quality of the image^37^ did not correlate with the GEDI-CNN accuracy, suggesting the GEDI-CNN is capable of making quality predictions on out-of-focus images (Supplemental Figure 2).

**Figure 2.**
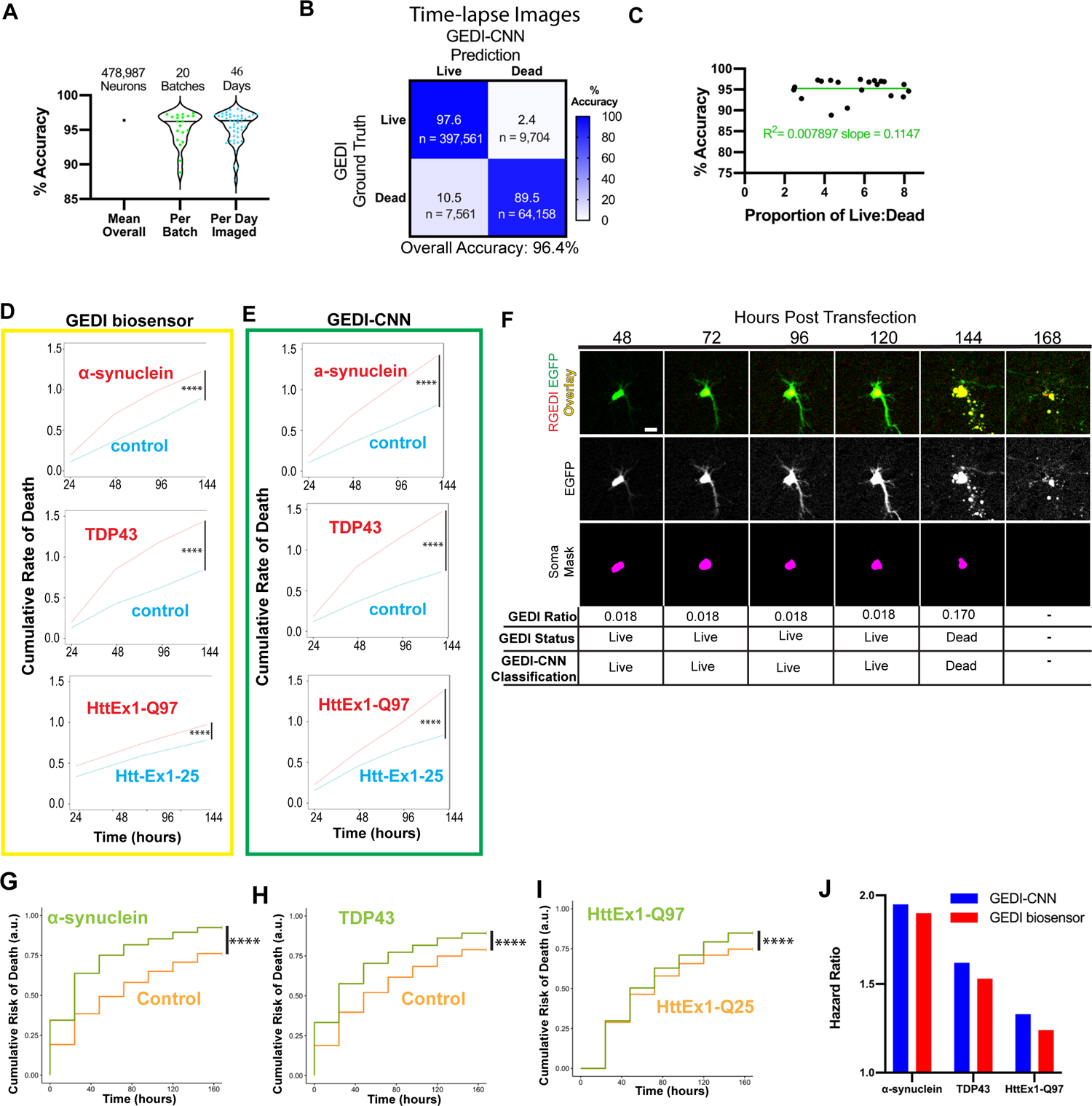
Application of GEDI-CNN to time lapse, single cell imaging of neurons. A) Accuracy of GEDI-CNN testing on time-lapse imaging across the entire data set, per batch (representing biological variation between primary neuron preps), and per imaging date (representing technical variability of the microscope over time). Black horizonal line represents mean. B) Confusion matrix comparing GEDI-CNN predictions to ground truth (GEDI biosensor) with percentage (top) and neuron count (bottom). C) Percent accuracy as a function of the proportion of live to dead neurons in data set. Green line represents linear regression fit to data. D, E) Cumulative rate of death per well over time for *α*-synuclein-expressing versus RGEDI-P2a-EGFP (control) neurons (top), TDP43-expressing versus control (middle) neurons, or HttEx1-Q97 (bottom) versus HttEx1-Q25 neurons using GEDI biosensor (D) or GEDI-CNN (E) (n= 48 wells each condition). F) Representative time lapse images of a dying single neuron with correct classification showing the overlay (top) (EGFP=green, RGEDI=red), the free EGFP morphology signal (middle) used by GEDI-CNN for classification, and the binary mask of automatically segmented object (bottom) used for quantification of GEDI ratio. By 168 hours post transfection, free EGFP within dead neurons has sufficiently degraded for an object not to be detected. G) Cumulative risk-of-death of α-synuclein-expressing versus RGEDI-P2a-control neurons (HR=1.95, **** p<2e-16), (H) TDP43 versus RGEDI-P2a-control neurons (HR=1.62, **** p<2e-16), and (I) HttEx1-Q97 versus HttEx1-Q25 neurons (HR=1.33, *** p<2.11e-13). J) Comparison of the hazard ratio using GEDI-CNN (top) versus GEDI biosensor^31^ (bottom). Scale = 20μm.

We next tested whether a GEDI-CNN could detect cell death in longitudinally-imaged rat primary cortical neuron models of neurodegenerative disease. Imaging with the GEDI biosensor previously showed that protein overexpression models of Huntington’s Disease (HD) (pGW1:HttEx1-Q97)^32^, Parkinson’s Disease (PD) (pCAGGs:α-synuclein)^38^, and amyotrophic lateral sclerosis (ALS) or frontotemporal dementia (FTD)(pGW1:TDP43)^26^ have high rates of neuronal death^31^. We therefore compared the cumulative rate of death per well (dead neurons/total neurons) as Figure 2 estimated by the GEDI biosensor versus the GEDI-CNN. We began by quantifying cumulative rate of death per well, which does not require tracking each neuron in the sample, simplifying the comparison between the GEDI biosensor and the GEDI-CNN predictions. From the cumulative rate of death, we derived a hazard ratio comparing each disease model with its control using a linear mixed effects model (Figure 2D, E). For each disease model, the GEDI biosensor and the GEDI-CNN detected similar increases in neuronal death in the overexpression lines compared to the control lines, confirming the GEDI-CNN can be used to automatically and quickly assess the amount of neuronal death in live imaging data (Figure 2D, E). Furthermore, the high accuracy of the GEDI-CNN indicates that there is enough information in the morphological changes of a neuron alone to accurately predict reduced survival within a neuronal disease model with nearly equivalent performance as the GEDI biosensor.

Evaluation of the amount of death in a cross-section of cells per well over time can be an effective tool for tracking toxicity within a biological sample, but it cannot resolve transient biological changes within the sample due to changes in the total cell number or heterogeneity within the culture. Changes in cell number are treated as technical variation, which reduces the overall sensitivity of the system and can mask transient changes within the culture. In contrast, single-cell longitudinal analysis has proven advantages over conventional imaging approaches because the live/dead state of each cell, the total cell number, and rare or transient changes in cells can be quantitatively linked to their fate^3^. In combination with a suite of statistical tools commonly used for clinical trials called survival analysis^39^ and Cox proportional hazard (CPH) analysis^40^, the analysis of time of death in single cells can provide 100–1000 times more sensitivity than the analysis of single snapshots in time^3^. Additionally, single-cell approaches can facilitate co-variate analyses of the factors that predict neuronal death ^28, 29, 41–43^. Yet, each of these analyses is limited by an inability to acutely differentiate live and dead cells^27, 31^.

We performed automated single-neuron tracking on HD, PD, and ALS/FTD model datasets while quantifying the timing and amount of death detected with GEDI biosensor or GEDI-CNN (Figure 2F). With accurate single-cell tracking and live/dead classification in place, we applied CPH analysis to calculate the relative risk of death for each model, which closely recapitulated previously reported hazard ratios (Figures 2G– J)^31^, and mirrored the cumulative rate of death analysis. These data indicate the GEDI-CNN, trained with GEDI biosensor data or GEDI-CNN from the first time point of live imaging alone, can accurately predict death in later timepoints using the morphology signal alone, which can be used to identify neurodegenerative phenotypes in longitudinal culture models.

### GEDI-CNN outperforms human curation

The high accuracy with which the GEDI-CNN classifies dead cells in imaging time series suggested that it could also be an improvement over standard human curation. To test this hypothesis, we manually curated 3,000 images containing GEDI biosensor that were not included in model training. Four trained human curators classified the same set of neurons as live or dead using the EGFP morphology signature by interacting with a custom-written ImageJ script that displayed each image and kept track of each input and the time each curator took to accomplish the task. The GEDI-CNN Figure 3

**Figure 3.**
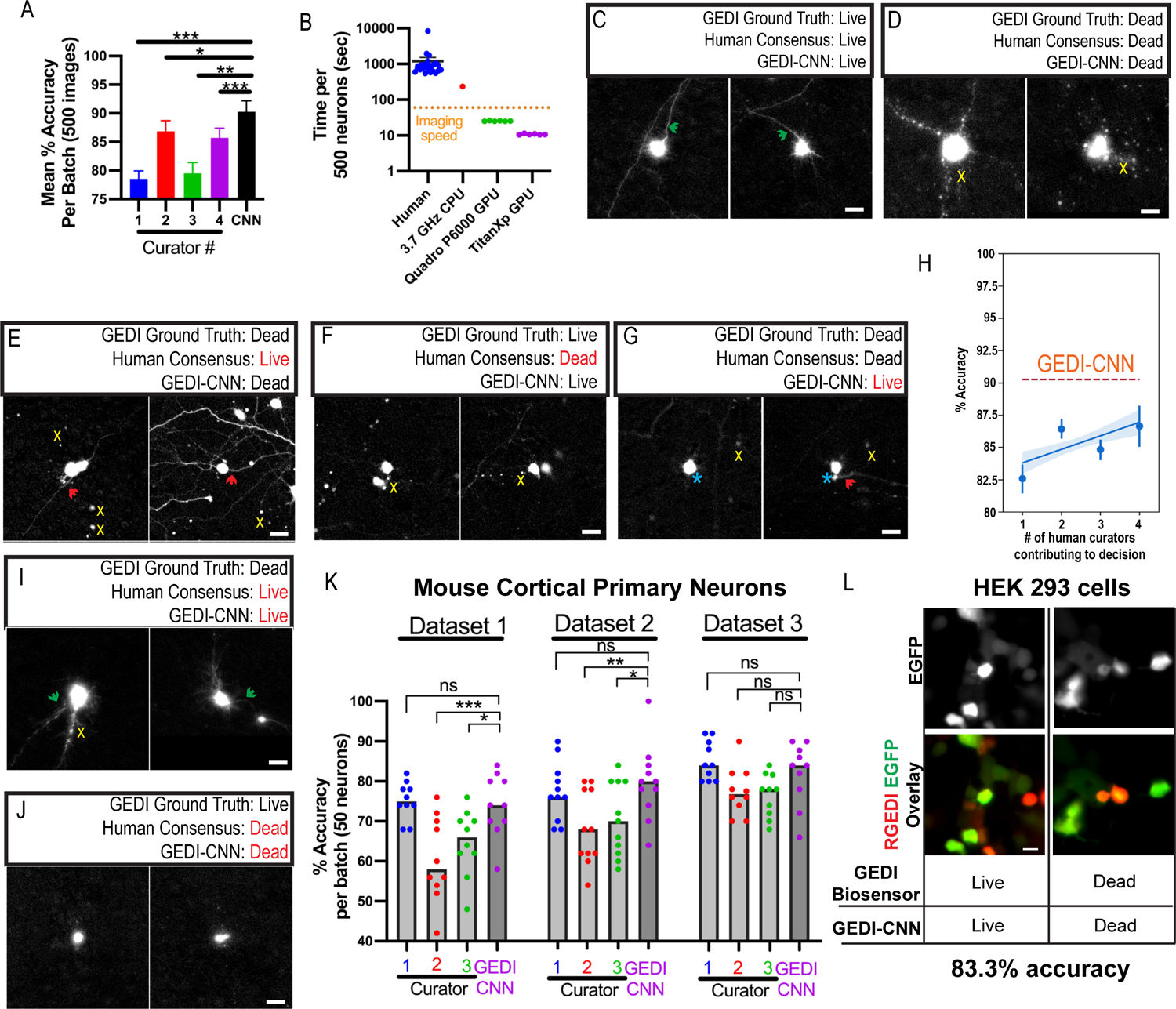
GEDI-CNN has super-human accuracy and speed compared to human curation at live/dead classification. A) Mean classification accuracy per batch of GEDI-CNN and four human curators across four batches of 500 images using GEDI biosensor data as ground truth (+/- SEM). B) Quantification of the speed of image curation by human curators versus GEDI-CNN running on a CPU or GPU. Dotted line indicates average imaging speed (+/- SEM). C–D) Representative cropped EGFP morphology images in which GEDI biosensor ground truth, a consensus of human curators, and GEDI-CNN classify neurons as live (C) or dead (D). Green arrows indicate presence of neurites from central neuron, yellow x’s indicate peripheral debris near central neuron. E–F) Examples of neurons in which GEDI ground truth and GEDI-CNN classified a neuron as dead whereas the human curator consensus was live (E), or GEDI ground truth and GEDI-CNN classified a neuron as live whereas the human curator consensus was dead (F). Red arrows indicate neurites that may belong to a neuron other than the central neuron. Yellow x’s indicate peripheral debris near the central neuron. G) Examples of neurons in which GEDI-CNN indicates the cell is live while GEDI ground truth and human consensus indicate that the cell was dead.

Turquoise asterisk indicates a blurry fluorescence from the central neuron, yellow x’s indicate peripheral debris near the central neuron. H) Accuracy achieved by human consensus based on the mean decision across increasing number of curators contributing to decision^69^. Blue line represents linear regression and shadow represents 95% CI of regression. Number of humans required for consensus accuracy to achieve GEDI-CNN accuracy. I) Examples of neurons classified as dead by GEDI ground truth and live by human consensus and the GEDI-CNN classification. Green arrows indicate presence of neurites from central neuron, yellow x’s indicate peripheral debris near central neuron. J) Examples of neurons classified as live by GEDI ground truth and dead by human consensus and GEDI-CNN classification. K) Mean classification accuracy of GEDI-CNN in comparison to human curators across three mouse primary cortical neuron datasets. Scale bar = 20μm. L) Representative cropped EGFP (top) and overlaid RGEDI and EGFP images (bottom) of HEK293 cells. Cell centered on left is live by both GEDI biosensor and GEDI-CNN. Cell centered on right panels is dead by both GEDI biosensor and GEDI-CNN.

was 90.3% correct in detecting death across six balanced batches of 500 images each, significantly higher than each human curator individually (curator range 79-86%) Figure 3A). In addition to accurately detecting cell death, the GEDI-CNN was two orders of magnitude faster than human curators at rendering its decisions on our hardware; for the first time, the rate of data analysis was faster than the rate of data acquisition (imaging) (Figure 3B). Images correctly classified live by the GEDI-CNN passed classical curation standards such as the presence of clear neurites in live neurons (Figure 3C). Neurons classified as dead showed patterns of apoptosis such as pyknotic rounding-up of the cell, retraction of neurites, and plasma membrane blebbing in dead neurons^44^(Figure 3D). These data suggest the GEDI-CNN learns to detect death using features in neuronal morphology that have been previously recognized to track with cell death.

Images for which the consensus of the human curators differed from that of the GEDI-CNN are of interest in finding examples of morphology related to death that are imperceptible to humans, and instances where the GEDI-CNN but not humans systematically misclassified death. For example, neurites were typically present in images where no human curator detected death, but the GEDI-CNN did (Figure 3E). This suggests that human curators may be overly reliant on the presence of contiguous neurites to make live or dead classifications. In cases where humans incorrectly labeled a neuron as dead while the GEDI-CNN correctly indicated a live neuron, images were often slightly out of focus, lacked distinguishable neurites, or contained nearby debris and ambiguous morphological features such as a rounded soma that are sometimes though not always indicative of death (Figure 3F). Similarly, in cases where the GEDI-CNN but not humans incorrectly classified dead neurons as alive, the neurons typically lacked distinguished neurites, contained ambiguous morphological features, and were surrounded by debris whose provenance was not clear (Figure 3G). We recorded no instances of live neurons that were correctly classified by humans, but incorrectly classified by the GEDI-CNN. These data suggest that the GEDI-CNN correctly classifies neurons with super-human accuracy, in part because it is robust to many imaging nuisances that limit human performance, which would explain how the accuracy of GEDI-CNN exceeded accuracy of curators, taken either individually or as an ensemble of curators (Figure 3H).

The curator consensus and the GEDI-CNN were still both incorrect in over 9% of the curated images, suggesting there are some images in which the live or dead state cannot be determined by the EGFP morphology alone. This could be in part because the level of Ca^2+^ within the neuron detected by GEDI is a better indicator of death than EGFP morphology. Close examination of images in which curation and GEDI-CNN were incorrect showed ambiguity in common features used to classify death, such as the combination of intact neurites and debris (Figure 3I), or lack of neurites and rounded soma (Figure 3J), suggesting the morphology of the image was not fully indicative of its viability. These data suggest that GEDI-CNN can exceed accuracy of human curation, but it is also limited by the information present in the EGFP morphology image.

CNNs often fail to translate across different imaging contexts, so we next examined whether GEDI-CNN accurately classifies within datasets collected with different imaging parameters and biology. To probe how well GEDI-CNN translates to neurons from other species, we prepared plates of mouse cortical primary neurons expressing the GEDI biosensor and imaged them with parameters similar though not identical to those used on the rat primary cortical neurons used for initial training and testing. The GEDI-CNN showed human-level accuracy across each mouse primary neuron dataset (Figure 3K, Supplemental Figure 3). While the GEDI-CNN significantly outperformed some human curators in some datasets, it did not have a consistent edge in accuracy over all human curators, suggesting some superhuman classification accuracy is lost when translating predictions to new biology or imaging conditions. To probe how far GEDI-CNN could translate, we next transfected immortalized human embryonic kidney (HEK) 293 cells, a cancer cell line, with RGEDI-P2a-EGFP, and exposed the cells to a cocktail of sodium azide and triton-X100 to induce death in the culture. HEK293 cells lack neurites and their morphology differs notably from the morphology of primary neurons, yet the GEDI-CNN classification accuracy was still significant—83.3% —in these cells (Figure 3L, Supplemental Figure 3). Overall, these data demonstrate that the GEDI-CNN provides superhuman speed and accuracy that generalizes across a diverse range of imaging, biological and technical conditions.

### GEDI-CNN uses membrane and nuclear signal to cue superhuman classifications

Our findings indicate that the GEDI-CNN joins a growing number of other examples that use CNNs to achieve superhuman speed and accuracy in biomedical image analysis^16, 45^. Nevertheless, a key limitation of these models is the difficulty in interpreting the visual features they rely on for their decisions. One popular technique for identifying the visual features that contribute to CNN decisions is Guided Gradient-weighted Class Activation Mapping (GradCAM)^46^. GradCAM produces a map of the importance of visual features for a given image by deriving a gradient of the CNN’s evidence for a selected class (*i.e.,* dead) that is masked and transformed for improved interpretability (Supplemental Figure 4A). Indeed, GradCAM has been found to identify visual features used by leading CNNs trained on object classification that closely align with those used by human observers to classify the same images^46^.

We generated GradCAM feature importance maps for both live and dead decisions for every image (Supplemental Figure 4B–D). We found that feature importance maps corresponding to the GEDI-CNN’s ultimate decision placed emphasis on cell body contours and neurites (Figure 4 A, B). GradCAM importance maps highlighted the central segmented soma rather than peripheral fluorescence signal that might have come from other neurons within the cropped field of view. These importance maps also did not indicate that regions without fluorescence were important for visual decisions (Supplemental Figure 4E–H). Thus, the GEDI-CNN learned to base its classification on the neuron in the center of the image rather than background nuisances or other neurons not centered in the image. Indeed, the backbone architecture of the GEDI-CNN— a VGG-16— is uniquely suited for learning this center bias due to its multi-layer, fully connected readout, which has distinct synapses for every spatial location and feature in the final convolutional layer of the model. While this bias is often not helpful in natural image classification, and has been replaced with spatially global average pooling in more recent architectures like residual networks (ResNets)^47^, it is particularly well-suited for our imaging pipeline, which automatically centers the target neuron in every image.

**Figure 4.**
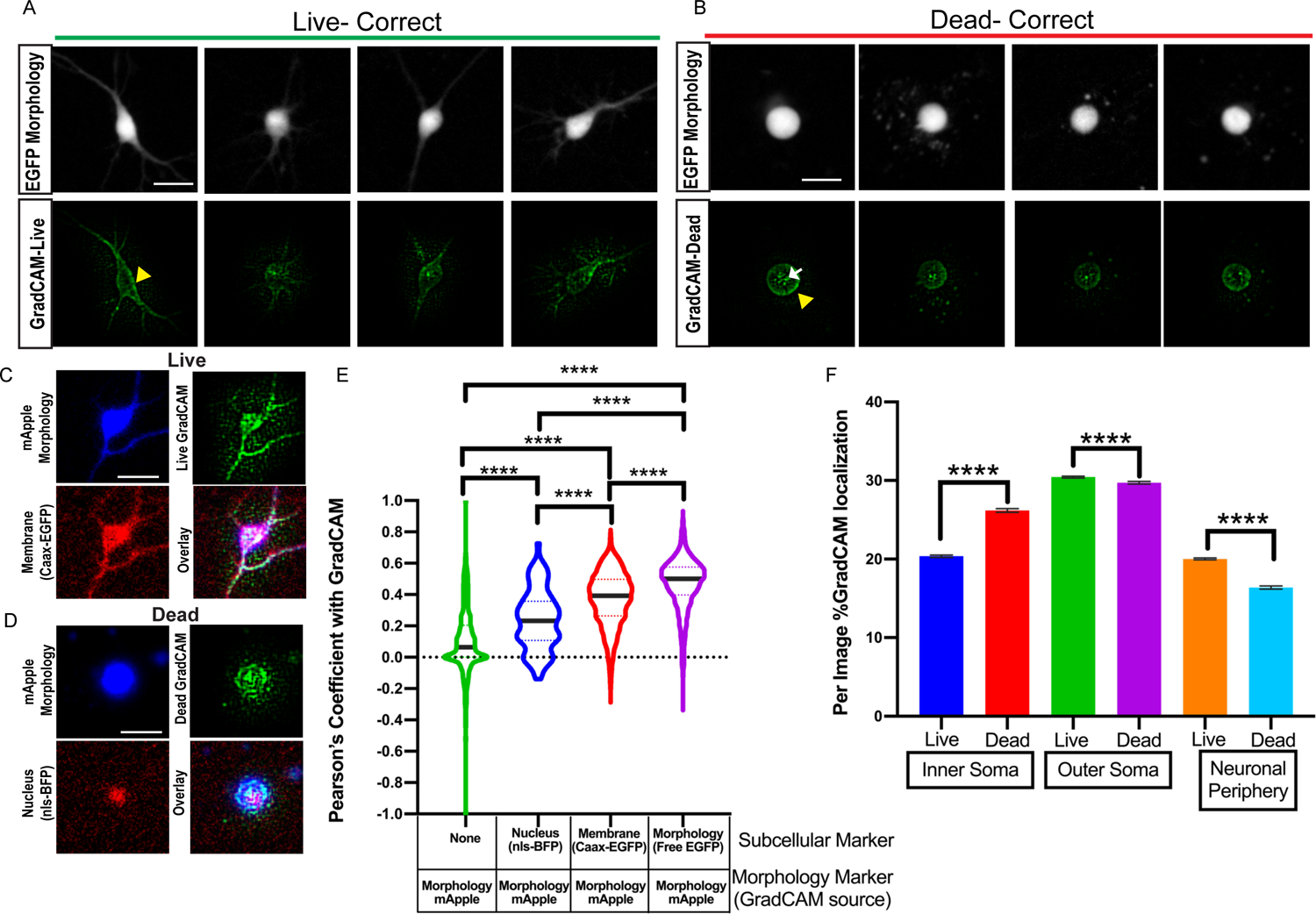
GEDI-CNN learns to use membrane and nuclear signal to classify death with superhuman accuracy. A–B) Free EGFP fluorescence of neurons (top) and GEDI-CNN GradCAM signal (bottom) associated with correct live (A) and dead (B) classification. Yellow arrowhead indicates GradCAM signal corresponding to the neuron membrane. White arrow indicates GradCAM signal within the soma of the neuron. C) Cropped image of a neuron classified as live by GEDI-CNN that is co-expressing free mRuby (top-left, blue) and CAAX-EGFP (bottom left, red), with the corresponding GradCAM image (green, top right), and the three-color overlay (bottom right). D) Cropped image of a neuron classified as dead by GEDI-CNN co-expressing free mRuby (top-left, blue) and nls-BFP (bottom left, red), with the corresponding GradCAM image (green, top right), and the three-color overlay (bottom right). E) Violin plot of quantification of the Pearson Coefficient after Costes’s automatic thresholding between GradCAM signal and background signal (n= 3083), nls-BFP (n = 86), CAAX-EGFP (n = 3151), and free mRuby (n=5404, **** = p <0.0001). F) Per image percentage of GradCAM signal in the inner soma and outer soma across live and dead neurons. Scale bar = 20μm.

We next looked to identify neuronal morphology that GEDI-CNNs deemed important for detecting dead neurons. The two consensus criteria under which a solitary cell can be regarded as dead according to current expert opinion are either the loss of integrity of the plasma membrane, or the disintegration of the nucleus^48^. We tested whether the GEDI-CNN had learned to follow the same criteria by performing subcellular co-localization of the GradCAM feature maps. Neurons were co-transfected with free mApple to mark neuronal morphology, and either nuclear targeted BFP (nls-BFP) or membrane targeted EGFP (Caax-EGFP). Pearson correlation between biomarker-labeled images and GradCAM feature maps were used to measure the overlap between the two^49^ (Figure 4C–E). GradCAM signal significantly co-localized with Caax-EGFP, and to a lesser extent nls-BFP fluorescence. To further validate the importance of nuclear and membrane regions for live/dead classification, we devised a heuristic segmentation strategy that further subdivides the neuronal morphology. We subdivided each masked soma into inner and outer regions (Supplemental Figure 5).

Signal outside of the soma was also separated into neuronal periphery (neurites and other neurons within the image crop) signal versus regions with no fluorescent signal. We found that signal within the inner soma was enriched for nls-BFP, whereas signals within the outer soma and neuronal periphery were enriched for membrane-bound EGFP (Caax-EGFP). The importance of the inner soma region to the ultimate decision classification was increased in neurons classified as dead versus those classified as alive, according to GradCAM feature importance analysis. In contrast, GradCAM feature importance decreased in the outer soma and neuronal periphery in live-classified neurons compared to dead-classified neurons (Figure 4F). These co-localizations Figure 5 indicate that the GEDI-CNN uses opposing weights for features near the nucleus versus the neurites and membrane in generating live/dead classification.

**Figure 5.**
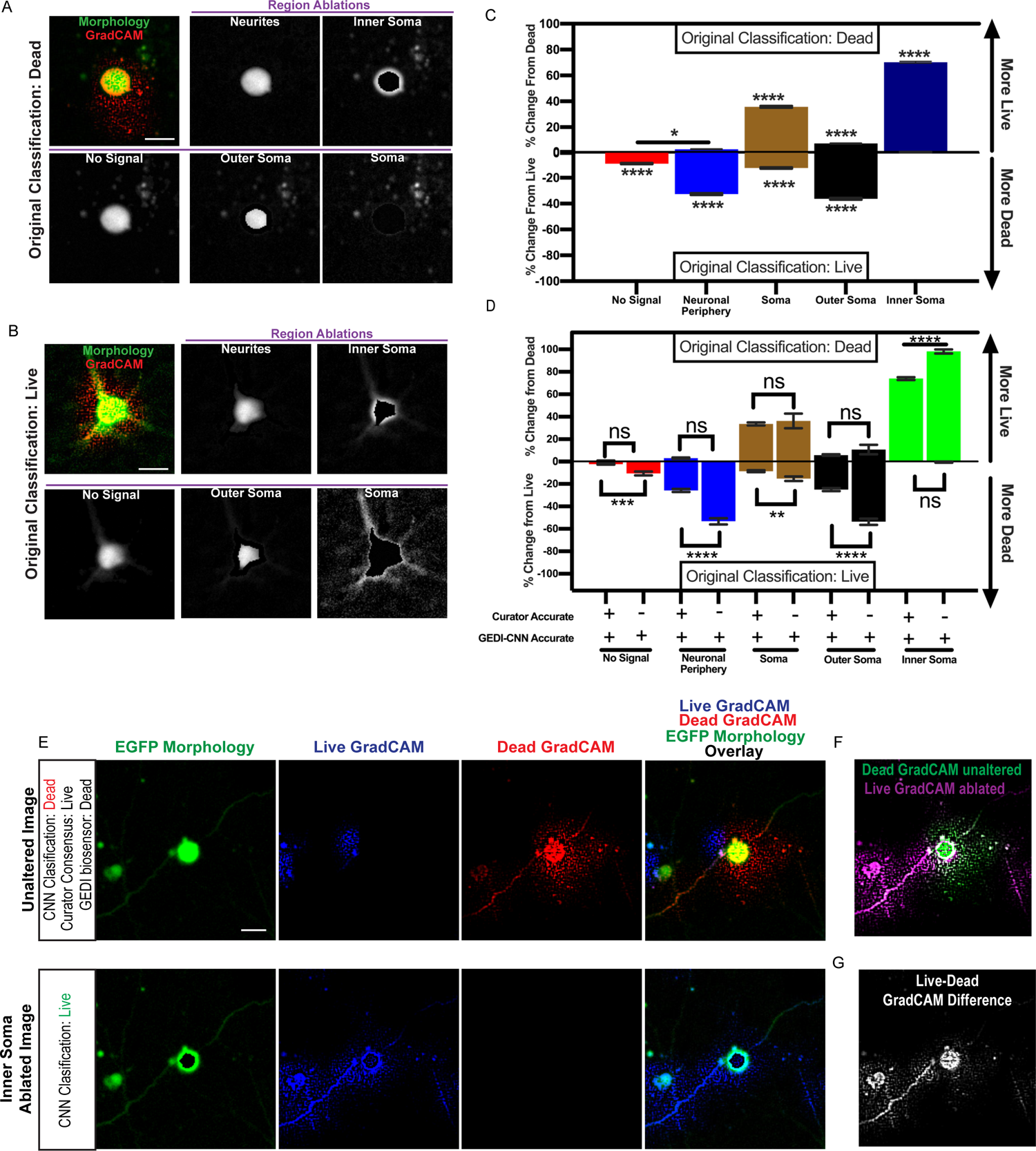
Inner soma signal important for dead classification, and membrane and neurite signals important for live classifications by GEDI-CNN A–B) (Top left) Overlay of free EGFP morphology (green) and GradCAM signal (red) from a neuron classified as dead (A) or live (B) by GEDI-CNN and a series of automatically generated ablations of the neurites, inner soma, no signal, outer soma, and soma (left to right, top to bottom). C) Effects of signal ablations on GEDI-CNN classification by region. Top box contains quantification per region of mean % change in classification of neurons originally classified as dead to live after ablation (n= 4619). Bottom box contains quantification of neurons originally classified as live (n=5122). Comparisons of all columns are significant (Tukey’s post hoc with correction for multiple comparisons ****= p<0.0001, comparison indicated * = p<0.05). D) Comparison of the effects of ablating signal from different subcellular regions on GEDI-CNN classification accuracy of neurons that were originally correctly classified by GEDI-CNN but incorrectly classified by at least two human curators (Tukey’s post hoc with correction for multiple comparisons, **** = p<0.0001, * = p<0.05). E) (Top) Representative example of EGFP morphology image of a dead neuron classified live by a consensus of four human curators and dead by a GEDI-CNN with its associated live and dead GradCAM signals. (Bottom) Inner soma-ablated EGFP image that changed GEDI-CNN classification from dead to live, and its associated live and dead GradCAM signals. F) Overlay of the dead GradCAM signal from the dead-classified, unaltered image and the live GradCAM signal from the live-classified inner soma-ablated image. G) Difference between dead GradCAM signal from the unaltered image and the live GradCAM signal from the inner soma-ablated image. Scale bar = 20μm.

GradCAM signal correlations suggested the nuclear/inner soma region and membrane/outer soma/peripheral neurite regions are important for live/dead classification, but they did not establish causality in the relationship of those features to the final classification. To validate the relationships, we measured GEDI-CNN classification performance as we systematically ablated pixels it deemed important for detecting death. We used heuristic subcellular segmentations to generate a panel of ablation images in which signal within each region of interest was replaced with noise, and we used these images to probe the decision making of the GEDI-CNN (Figure 5A– B). Ablations of the neurites and outer soma had the greatest effect in changing classifications of live neurons to dead, while ablations of the inner soma had the least effect (Figure 5C bottom). In contrast, ablation of the soma, particularly the inner soma, caused the GEDI-CNN to change dead classifications to live substantially more often than ablations of no-signal, neurites and outer soma (Figure 5C top). Although the inner soma and outer soma regions were the most sensitive ablations for switching classification to dead or live respectively, they represented the smallest areas of ablated pixels, suggesting the correct classification is based on a very specific subcellular morphological change (Supplemental Figure 6). These data confirm that regions enriched in membrane signal and in nuclear signal are critical for GEDI-CNN live-dead classification, though in opposite ways.

We next investigated if the source of the super-human ability of the GEDI-CNN to classify neurons as live or dead was a result of the CNN’s ability to see patterns in images that humans do not easily see within these identified regions. To this end, we analyzed the impact of regional ablations in images that the GEDI-CNN classified correctly but that more than a single curator misclassified (Figure 5D). Strikingly, almost all dead neurons that GEDI-CNN correctly classified but human curators incorrectly classified required the inner soma signal, suggesting that the inner soma signal underlies GEDI-CNN’s superhuman classification ability. GradCAM signal showed a shift from within the inner soma to the membrane and neurites after ablation (Figure 5E–G). These data show that the GEDI-CNN used the nuclear/inner soma and neurite/membrane morphology to gain superhuman accuracy at live or dead classification of neurons.

### Zero-shot transfer application of GEDI-CNN to human iPSC-derived motor-neurons

We next asked whether GEDI-CNN could detect death in a neuronal model of ALS derived from human induced pluripotent stem cells (iPSCs). Neurons derived from iPSCs (i-neurons) maintain the genetic information of the patients from whom they are derived, facilitating modeling of neurodegenerative^50^, neurological^51^, and neurodevelopmental diseases^52^ in which cell death can play a critical role. iPSC motor neurons (iMNs) generated from the fibroblasts of a single patient carrying the SOD1 D90A mutation have been previously shown to model key ALS-associated pathologies, and display a defect in neuronal survival compared to neurons derived from control fibroblasts^31, 53, 54^. However, iPSC lines derived from patient fibroblasts with less common SOD1 mutations associated with ALS, I113T^55–57^ and H44R^58^, have not been previously characterized for survival deficits. Whether reduced survival in culture is a common characteristic of all iMNs with SOD1 mutation remains unknown. To test if the GEDI-CNN can detect reduced survival in human iPSC-derived neurons, we differentiated iPSCs from SOD1^I113T^ and SOD1^H44R^ ALS patients and healthy volunteers into iMNs, transfected neurons with RGEDI-P2a-EGFP, and imaged them longitudinally (Figure 6A–C). Overall classification accuracy based on morphology alone of i-neurons was 86.1% (0.68 AUC), with a markedly higher accuracy for live neurons (91.2%) than dead ones (41.4%) (Figure 6D). Across balanced randomized batches of data, GEDI-CNN averaged only 70% accuracy, and was not significantly different from human curation accuracy, indicating that death is difficult to discern from the EGFP morphology alone in this dataset (Figure 6E). A potential explanation for this difficulty is that iMNs often underwent a prolonged period of degeneration characterized by neurite retraction before a positive GEDI signal was observed (Figure 6C), a phenomenon not observed in mouse primary cortical neurons. This problem would make death difficult to discern from EGFP morphology alone in this dataset.

**Figure 6.**
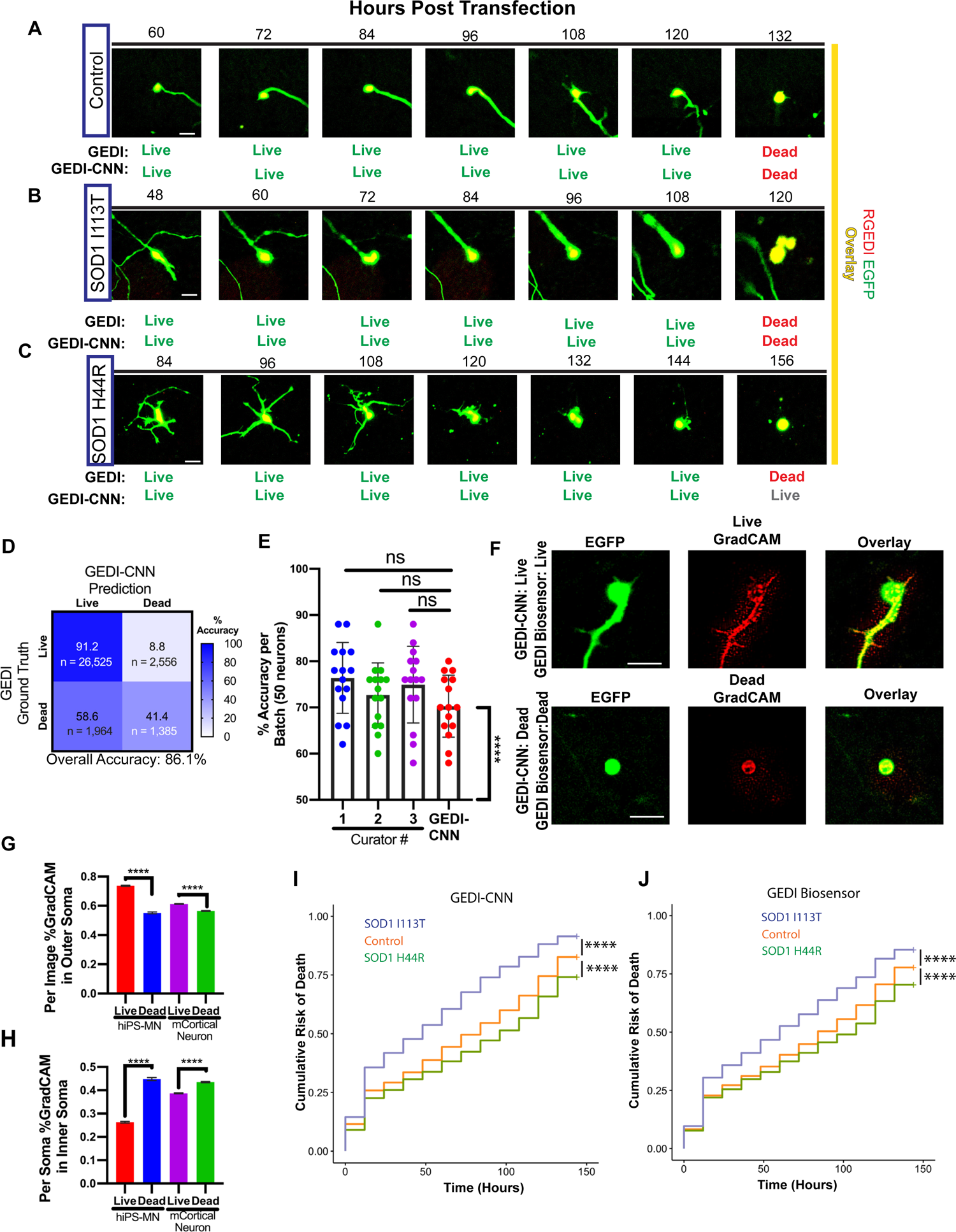
GEDI-CNN classifications translate to studies of neurodegeneration in human iPSC-derived motor neurons A–C) Representative longitudinal time lapse overlay images RGEDI (red) and EGFP (yellow) in iPSC-derived motor neurons derived from a healthy control patient (A), an ALS patient with SOD1^I113T^ (B), or an ALS patient with SOD1^H44R^ (C). Comparison of live/dead classifications derived from the GEDI biosensor and GEDI-CNN below each image. D) Confusion matrix of live/dead classification accuracy of GEDI-CNN on iPSC-derived motor neurons. E) Mean classification accuracy of GEDI-CNN and human curators on randomized 50% live:dead balanced batches of data (ns = not significant, ANOVA with Tukey’s multiple comparisons). Each curator and the GEDI-CNN showed classification accuracy significantly above chance (**** = p<0.0001 Wilcoxon signed-rank difference from 50%). F) Representative images of free EGFP morphology expression (Left), GradCAM signal (Middle), and overlay of GradCAM and EGFP (Right) of iPSC-derived motor neurons classified correctly as live (top) and dead (bottom) by GEDI-CNN. G–H) GradCAM signal in the soma shifts from outer soma in live neurons (G) to inner soma in dead neurons (H) in both human iPSC-derived motor neurons (hiPS-MN) and mouse primary cortical neurons (mCortical Neuron). (I) Cumulative risk-of-from death of SOD1^I113T^ (HR=1.52, ****, p<0.0001) and SOD1^H44R^ (HR=0.79, ****, p<0.0001) neurons derived from GEDI-CNN. (J) Cumulative risk-of-from death of SOD1^I113T^ (HR=1.35, ****, p<0.0001) and SOD1^H44R^ (HR=0.83, ****, p<0.0001) neurons using GEDI biosensor. Scale bar = 20μm.

GradCAM signal from correctly classified live and dead neurons defined the contours of neurons of iMNs (Figure 6F) and showed similar signal changes in response to neuronal death as observed in mouse cortical neurons (compare Figure 6 G, H to Figure 4). Single-cell KM and CPH analysis of SOD1^I113T^ and SOD1^H44R^ neurons yielded similar results with the GEDI-CNN classification and the GEDI biosensor classification. In both cases, SOD1^I113T^ neurons showed reduced survival compared to control neurons while SOD1^H44R^ neurons showed significantly increased survival (Figure 6 I, J). Human curation using neuronal morphology showed the same trends for (Supplemental Figure 7). The hazard curves and HR obtained with GEDI-CNN for SOD1^D90A^ were also similar to previously reported GEDI biosensor data (Supplemental Figure 8)^31^. These data demonstrate that a reduced survival phenotype of iMNs generated from patients with ALS is consistently detected in some cell lines (SOD1^D90A^, here,^31, 53, 54^), but not detected in other others (SOD1^H44R^), suggesting that higher-powered studies are necessary to overcome inherent study variability and attribute phenotypes found in iMNs to underlying genetic factors. Overall, our data show that the GEDI-CNN can be deployed to make human-level zero-shot predictions of GEDI ground truth on an entirely different cell type than the one on which it was trained.

## Discussion

Microscopy is frequently hindered by the rate and accuracy of image analysis. Here, we developed a novel strategy that leverages physiological biomarkers to train computer vision models and automate image analysis. Using signal from GEDI, a sensitive detector of Ca++ levels presaging cell death (ref), we trained a CNN that detects cell death with superhuman accuracy and performs across a variety of tissues and imaging configurations. Importantly, we demonstrate that the GEDI-CNN learned to detect cell death by locating fundamental morphological features known to be related to cell death, despite receiving no explicit supervision to focus on those features.

Additionally, we demonstrate that in combination with automated microscopy, we can accelerate the analysis of large datasets to rates faster than the rate of image acquisition, enabling highly sensitive single-cell analyses at large scale for the first time.

Using this strategy, we identified distinct survival phenotypes in iPSC-derived neurons from ALS patients with different mutations in SOD1. Our results demonstrate that the GEDI-CNN can improve the precision of experimental assays designed to uncover the mechanisms of neurodegeneration, and that the BO-CNN strategy can be used to accelerate discovery in the field of live-cell microscopy.

GEDI-CNN live/dead classification predictions were significantly above-chance across every cell type we tested on (Supplemental Figure 3). Given the diversity of cell types tested, this indicates GEDI-CNN learned to locate morphology features that are fundamental to death across biology. Using a combination of GradCAM and pixel ablation simulations, we show that the predictions of “live” and “dead” are sourced from different subcellular compartments such as the plasma membrane and the nucleus.

Altered plasma membrane and retracted neurites are prominent features in images of neuronal morphology that are well known to correlate with vitality^59^, and are likely easily appreciated by human curators. However, EGFP morphology signal in the region of the nucleus has not been previously appreciated as an indicator of vitality. While rupture of the nuclear envelop is well-recognized as an indication of cell vitality^48^, it is not clear how free EGFP signal that is both inside and outside of the nuclear envelope could be indicating nuclear envelope rupture, or if another cell death-associated phenomenon is responsible for this signal. Either way, EGFP morphology signal in the nuclear region of cell represents a previously unrecognized feature that correlates with cell death, and one that helps the GEDI-CNN achieve superhuman classification accuracy. The ability to harness the superhuman pattern recognition abilities of BO-CNN models represents a powerful new direction for the discovery of novel biological phenomena and underscores the importance of understanding the visual basis of classification models to interpret biology.

Our demonstration that the GEDI-CNN can automate live-imaging studies of cellular models of neurodegenerative disease has tremendous implications for accelerating research in this field. Neuronal death is a nearly-ubiquitously used phenotypic readout in studies of neurodegeneration^27^. We demonstrate the use of the GEDI-CNN for large-scale studies of neurons using accurate cell death as a readout for the first time, and its potential for high-throughput screening purposes. Similarly, in large-scale studies or high-throughput screens of neurons in which cell death is not the primary phenotype, the GEDI-CNN can accurately account for cell death or be used to filter out dead objects that could contaminate other analyses. Employing the GEDI-CNN to make predictions and map GradCAM on morphology images of neurons is free, open source, simple to execute, and does not require specialized hardware. In addition to facilitating high-throughput screening, which can dramatically improve statistical power and sensitivity of analyses^3^, generally accounting for cell death in microscopy experiments should help control experimental noise.

A rapid rate of analysis is particularly necessary where high rates of heterogeneity and biological diversity are present. Studies using human iPSC-derived neurons have begun grappling with the importance of scale for identifying and studying biological phenomena in the incredibly diverse human population^60–62^. This is well-illustrated in our data applying GEDI and the GEDI-CNN to iMNs from patients with SOD1 mutations. Multiple studies of iPSC-derived motor neurons from patients with SOD1 mutations linked to ALS, such as A4V and D90A^31, 53^, have shown reduced survival phenotypes, and we observed a similar phenotype in cells harboring the I113TE and D90A mutations. However, the current study shows that the H44R mutation improves the survival of iPSC-derived neurons compared to control, indicating a cell death phenotype in iMNs from ALS patients under these conditions is not consistent under these conditions. There is high variability across different iPSC differentiation protocols, and it remains to be determined how much variability between iPSC lines contributes to previous underpowered reports of phenotypes associated with different lines. Ultimately the only way to overcome issues with iPSC line variability is to increase the amount of cell lines tested and standardize methods of measurements and analysis compatible with higher powered studies. As GEDI-CNN significantly reduces the time and resources required to do high-power studies and eliminates human bias and intra-observer variability in live/dead classification, GEDI-CNN live/dead classifications should become a useful tool in future studies attempting to link iMN phenotypes to their genotypes.

While GEDI-CNN model prediction accuracy was always above chance, the accuracy level was reduced in some datasets, suggesting differences in imaging parameters and/or biology may limit its ability to generalize from one data set to the next (Supplemental Figure 3). To offset reduced accuracy, new GEDI-CNN models can be transferred, or “primed”, with small datasets that better represent the qualities of a particular experiment or microscope setup. These datasets can be generated by imaging the GEDI biosensor on a particular microscope platform and training a model according to the biomarker optimized (BO-CNN) pipeline we outline here. Nevertheless, on some datasets we observed that both human and GEDI-CNN classification accuracy and GEDI-CNN accuracy were reduced, suggesting that in these cases morphology alone may be insufficient to accurately classify a cell as live or dead. Morphology of iMN datasets proved particularly difficult to interpret for both human curators and GEDI-CNN (Figure 6C). Interestingly, while classification accuracy for live neurons was consistently high across different cell types, accuracy of classification for dead neurons often reduced overall accuracy (Figure 2B, 6D), suggesting death classification by morphology is especially difficult. Because of this variability in how different biological systems reflect death, the use of the GEDI biosensor as a ground truth to evaluate GEDI-CNN model predictions as well as for training new models will continue as the gold standard for detecting cell death.

While GEDI-CNN represents the first proof of principle in using a live biosensor to train a BO-CNN, we believe similar strategies could be used to generate new BO-CNNs that could help parse mechanisms underlying other disease-related phenotypes in neurodegenerative disease models. Several features of the GEDI biosensor proved critical for success. First, the GEDI biosensor acts as a nearly binary demarcation of live or dead cell states^31^. Other biosensors with similar biological state-demarcation characteristics would make good candidates for this approach. However other strategies for automatically extracting a classification label, such as using image timing or applied perturbations as the classification label, could also be used as a ground truth in place of GEDI signal. Second, the GEDI signal is simple and robust enough to extract with currently-available image analysis techniques. However, the fluorescent proteins within the GEDI we used here (RGEDI and EGFP), are not the brightest in their respective classes, and many available biosensors could be used that have as much, if not more, signal. Third, the free EGFP morphology signal within GEDI is already well-known to indicate live or dead state, as it had been previously used as the gold-standard for longitudinal live-imaging studies of neurodegeneration^27, 32^. This gave us confidence that the patterned information within the EGFP morphology images would be sufficient to inform the model of the biological state of the neuron. While that information may not be available in all situations, there are now many examples where deep learning has uncovered new unexpected patterns in biological imaging, and the BO-CNN strategy could also be used in a similar way to uncover unexpected relationships between images and known biological classifications. As each of these properties can be found in other biosensors and biological problems, we are confident that other effective BO-CNNs can be developed using the same strategy we used to generate the GEDI-CNN. As the rate of microscopy imaging and scale has continued to accelerate in recent years from the commercialization of automated screening microscopes, we expect the importance of BO-CNN analyses to continue to increase, leading to new discoveries and therapeutic approaches to combat neurodegenerative disease.

## Methods

### Animals, Culturing, and Automated Time-lapse Imaging

All animal experiments complied with UCSF regulations. Primary mouse cortical neurons were prepared at embryonic days 20–21 as previously described^18^. Neurons were plated in a 96-well plate at 0.1 × 10^6^ cells per well and cultured in neurobasal growth medium with 100× GlutaMAX, Pen/Strep, and B27 supplement (NB medium).

Automated time-lapse imaging was performed with long time intervals (12–24 hours) in order to minimize phototoxicity while imaging, reduce data set size, and provide a protracted time window to monitor neurodegenerative processes. Imaging speed was assessed as the average time to image a 4×4 montage of EGFP-expressing neurons per well in a 96-well plate (6 seconds) divided by the average number of neurons transfected per well (50). HEK cell were plated in a 96-well plate at 0.01 × 10^6^ cells per well 24 hours before transfection. 0.4% sodium azide and 0.02% Triton-X100 (final concentration) were added to induce cell death. Images were captured 20 min after sodium azide and Triton-X100 application.

### Automated Imaging and Image Processing Pipeline

Quantification of GEDI ratios was performed as previously described^31^. In short, RGEDI and EGFP channel fluorescence images obtained by automated imaging were processed using custom scripts running within a custom-built image processing Galaxy bioinformatics cluster^63, 64^. The Galaxy cluster links together workflows of modules including background subtraction of median intensity of each image, montaging of imaging panels, fine-tuned alignment across time points of imaging, segmentation of individual neurons, and cropping image patches where the centroid of individual neurons is positioned at the center. Single-cell tracking was performed using a custom Voronoi tracking algorithm that extracts intensity and feature information from each neuron^31^. Survival analysis was performed by defining the last timepoint alive as the timepoint before the GEDI ratio of a longitudinally imaged neuron exceeds the empirically calculated GEDI threshold for death, the last timepoint a human curator tracking cells finds a neuron alive, or the timepoint before the GEDI-CNN classifies a neuron as dead. KM survival curves and CPH were calculated using custom scripts written in R, and survival functions were fit to these curves to derive cumulative survival and risk-of-death curves that describe the instantaneous risk of death for individual neurons as previously described^65^. Cumulative risk of death analysis was performed by quantification of cumulative mean of dead cells/total cells per well across timepoints.

Because fluorescent neuronal debris can remain in culture for up to 48 hours past the time point of death^31^, timepoints other than at 48 hour intervals were removed from the analysis to avoid double counting dead cells within the culture. A linear mixed-effects model implemented in R^66^ was fit to the cumulative rate of death, using “well” as a random effect and interaction between condition and elapsed hours as a fixed effect to derive hazard ratios. Corrections for multiple comparisons were made using Holm-Bonferroni method.

### Training GEDI-CNN

We began with a standard VGG16^36^, implemented in TensorFlow 1.4. The 96,171 images of rodent primary cortical neurons were separated into training and validation folds (79%/21%), normalized to [0, 1] using the maximum intensity observed in our microscopy pipeline, and converted to single precision. We adopted a transfer-learning procedure, in which we initialized early layers of the network with pretrained weights from natural image categorization (ImageNet^36^), and later layers of the network with random initializations. We empirically selected which layers to put at the end of the network (i.e., from moving from the readout towards the input) by training models with different amounts of these layers randomly initialized and recording their performance on the validation set. Models were trained on Nvidia Titan X GPUs for 20 epochs using the Adam optimizer^67^ to minimize class-weighted cross entropy between live/dead prediction and GEDI-derived labels. To improve model robustness, neuron images were augmented with random up/down/left right flips, cropping from 256×256 pixels to 224×224 pixels using randomly placed bounding boxes, and random rotations of 0°, 90°, 180, or 270°. These images, which were natively 1-channel intensity images were replicated into 3-channel images to match the number of channels expected by the VGG16. We selected the best performing weights according to validation loss for the experiments reported here. Experiment code can be found at https://github.com/finkbeiner-lab/GEDI-ORDER.

### GEDI-CNN GradCAM

Feature importance maps were derived using Guided GradCAM – a method for identifying image pixels that contribute to model decisions. Guided GradCAM is designed to control for visual noise that emerges from computing such feature attributions for visual decisions through very deep neural network models. After extracting these feature importance maps from our networks, which were the same height/width/number of channels as the input images, we visualized them by rescaling pixel values into unsigned 8-bit images, converting to greyscale intensity maps, then concatenating with a greyscale version of the input image to highlight morphology selected by the trained models to make their decisions.

### Image Ablations and Curation Tools

Curation was performed using a custom Fiji script that runs a graphical interface with a curator, displaying a blinded batch of cropped EGFP morphology images one at a time while prompting the curator to indicate whether the displayed neuron is live or dead with a keystroke (ImageCurator.ijm). Automated subcellular segmentations were generated and saved as regions of interest (ROIs) and measured within a custom Fiji Script (measurecell8.ijm). ROIs were used to generate a panel of ablations for each image using another custom Fiji script (cellAblations16bit.ijm). Pearson co-localization was measured using JACoP with Costes background randomization^49^ within a custom Fiji script (PearsonColocalization.ijm).

### iPSC Differentiation to MNs

NeuroLINCS iPSC lines derived from fibroblasts tissue from a healthy control individual or ALS patients with SOD1 H44R (CS04iALS-SOD1H44Rnxx) or I114T mutations (SOD1I114Tnxx) were obtained from the Cedars Sinai iPSC Core Repository. Healthy and SOD1 mutant iPSCs were found to be karyotypically normal, and were differentiated into MNs using a modified dual-SMAD inhibition protocol^68^ (http://neurolincs.org/pdf/diMN-protocol.pdf). iPSC-derived MNs were dissociated using trypsin (Thermo Fisher), embedded in diluted Matrigel (Corning) to limit cell motility, and plated onto Matrigel-coated 96-well plates. From day 20–35, the neurons underwent a medium change every 2–3 days. At day 32 of differentiation, neurons were transfected using lipofectamine, and imaged starting on day 33.

## AUTHOR CONTRIBUTIONS

JWL, DL and SF wrote the manuscript. Conceptualization by JWL, DL, SF, NC, AJ, and TS. Data analysis and statistics by JWL, DL, KS, and NC. Robotic microscopy, transfections, iPSC differentiation, and cell culture performed by JK, JL, NC, SW, and VO. GEDI-CNN code base developed by DL. GEDI-CNN training, GradCAM visualization, and Defocus-CNN evaluations performed by DL, JL, GR, and ZT. Custom Fiji scripts for analysis and curation of imaging experiments done by JWL and JL, and GR. All authors reviewed the manuscript.

## CODE AVAILABILITY

Code is available at https://github.com/finkbeiner-lab/GEDI-ORDER. Code for statistics and data preprocessing, is available upon request to the corresponding author. Access to and use of the code is subject to a non-exclusive, revocable, non-transferable, and limited right for the exclusive purpose of undertaking academic, governmental, or not-for-profit research. Use of the code or any part thereof for commercial or clinical purposes is strictly prohibited in the absence of a Commercial License Agreement from the J. David Gladstone Institutes.

## DATA AVAILABILITY

The data that support the findings of this study are available from the corresponding author upon reasonable request.

## ACKNOWLEDGEMENTS

This work was supported by grants from the NIH (U54 NS191046, R37 NS101996, RF1 AG058476, RF1 AG056151, RF1 AG058447, P01 AG054407, U01 MH115747), as well as support from the Koret Foundation Artificial Intelligence Program for Biomedical Research, the Taube/Koret Center for Neurodegenerative Disease Research, the Target ALS Foundation, the Amyotrophic Lateral Sclerosis Association Neuro Collaborative, a gift from Mike Frumkin, and the Department of Defense award W81XWH-13-ALSRP-TIA. Gladstone Institutes received support from a National Center for Research Resources Grant RR18928. We gratefully acknowledge the support of NVIDIA Corporation with the donation of the Titan Xp and Quadro P6000 GPUs used for this research. Kathryn Claiborn and Françoise Chanut provided editorial assistance, Kelley Nelson and Gayane Abramova administrative assistance, and David Cahill provided lab neuron curation, lab maintenance and organization. We would like to thank Reuben Thomas and the Gladstone Institutes bioinformatics core for assistance with statistical models.

